# Ammonia-oxidizing archaea release a suite of organic compounds potentially fueling prokaryotic heterotrophy in the ocean

**DOI:** 10.1101/558726

**Authors:** Barbara Bayer, Roberta L. Hansman, Meriel J. Bittner, Beatriz E. Noriega-Ortega, Jutta Niggemann, Thorsten Dittmar, Gerhard J. Herndl

**Affiliations:** Division of Bio-Oceanography, Department of Limnology and Bio-Oceanography, Center of Functional Ecology, University of Vienna, Vienna, 1090, Austria; International Atomic Energy Agency – Environment Laboratories, Radioecology Laboratory, 98000 Monaco, Monaco; ICBM-MPI Bridging Group for Marine Geochemistry, University of Oldenburg, 26129 Oldenburg, Germany; Department of Marine Microbiology and Biogeochemistry, NIOZ Royal Netherlands Institute for Sea Research, and Utrecht University, 1790 AB Den Burg, Texel, The Netherlands

## Abstract

Ammonia-oxidizing archaea (AOA) constitute a considerable fraction of microbial biomass in the global ocean, comprising 20-40% of the ocean’s prokaryotic plankton and thus play an important role in global nitrogen cycle. However, it remains enigmatic to what extent these chemolithoautotrophic archaea are releasing dissolved organic matter (DOM). A combination of targeted and untargeted metabolomics was used to characterise the exometabolomes of three model AOA strains of the *Nitrosopumilus* genus. Furthermore, we compared the composition of intra- and extracellular dissolved free amino acids (DFAA). Our results indicate that marine AOA exude a suite of organic compounds with potentially varying reactivity, dominated by nitrogen-containing compounds. A significant fraction of the released DOM consisted of labile compounds, which typically limit prokaryotic heterotrophic activity in open ocean waters, including amino acids, thymidine and B vitamins. In growing *Nitrosopumilus* cultures, hydrophobic amino acids were likely released as a result of passive diffusion corresponding to ammonia oxidation activity, while glycine was continuously released at high rates. Our results suggest that AOA release several ecologically and biochemically relevant metabolites, potentially fueling heterotrophic prokaryotes in the ocean.

## Introduction

Dissolved organic matter (DOM) is one of the Earth’s largest reactive carbon pools, similar in magnitude to atmospheric CO_2_ and of major significance for the global carbon cycle and climate^1,2^. DOM is highly complex, consisting of thousands of different organic molecules^3,4^ and comprising the most heterogeneous and dynamic pool of carbon in the oceans^5^. Its concentration and composition are affected by biotic processes such as photosynthesis and heterotrophic metabolism^6,7^, as well as by photochemical processes^8^. However, while the flux of carbon through this pool of molecules is mediated largely by microbial activity, the relationships between marine microbes and the molecules making up the DOM pool remain poorly characterized^9^.

DOM is released as a by-product of metabolically active microbes, but may also be released for nutrient acquisition and communication (in the forms of metal-binding ligands and quorum sensing chemicals, respectively)^6,10,11^, as well as upon predation or viral lysis^12^. The fraction of DOM exuded by phytoplankton is highly variable accounting for 5 - 70% of photosynthetically fixed carbon^6^. There are major gaps, however, in our understanding of the biogeochemical significance of the released compounds^13^. The composition of DOM released by heterotrophic bacteria is even less known than for phytoplankton, however, they do produce and release organic compounds for similar purposes as phytoplankton^5,6,14,15^.

Although the largest fraction of DOM remains uncharacterized with respect to molecular identity, phytoplankton exudates consist to a considerable extent of labile compounds such as carbohydrates (mono-, and polysaccharides), proteins and amino acids^16–18^. Hence, extracellular release of DOM by phytoplankton supports a major fraction of the labile carbon flux in the surface ocean, thereby fueling secondary production^18–21^. Labile DOM is defined as being consumed by microbes within hours to days of production but typically only accounts for a fraction of the bacterial C- and N-demand^22^. However, the majority of the oceanic DOM pool appears to be recalcitrant with lifetimes from years to millennia and its sources and sinks are largely unknown^23^.

Novel techniques based on electrospray ionization coupled to mass spectrometry allow for a more detailed characterization of the low molecular weight (LMW, < 1000 Da) DOM pool, which represents the major component of DOM and was not analytically accessible before^24^. The use of Fourier transform ion cyclotron resonance mass spectrometry (FT-ICR-MS) makes it possible to determine elemental formulae from highly accurate mass measurements alone^25^. These recent methodological advances provided new insights into the vast complexity of DOM secreted by phytoplankton and heterotrophic bacteria, suggesting that a major fraction of the released organic molecules is highly diverse and dependent on nutrient availability, the organism itself and its growth stage^26–29^. The DOM composition appears to be highly variable and microbes exude different compounds in the presence of other microbes^30^. Furthermore, auxotrophy (=the inability of an organism to produce a specific compound it requires) and the consequential exchange of a variety of metabolites and precursors between different microbes appear to be more widespread than previously assumed^31–34^.

Ammonia-oxidizing archaea (AOA) constitute a considerable fraction of microbial biomass in the global ocean, comprising 20-40% of the ocean’s prokaryotic plankton^35^. It is unknown to what extent these chemolithoautotrophic archaea are releasing DOM. If they do, it might be that a significant fraction of marine DOM is of archaeal origin. Furthermore, potential reciprocal interactions via DOM transformations between AOA and heterotrophic bacteria have been suggested previously^36^. Here, we investigated the exometabolomes of three model strains of marine AOA, *Nitrosopumilus adriaticus* NF5, *Nitrosopumilus piranensis* D3C and *Nitrosopumilus maritimus* SCM1, using untargeted and targeted metabolomics. While untargeted metabolomics aims at detecting and describing as much of the metabolome as possible, targeted metabolomics uses authentic standards to quantify a specific set of compounds^37^. Moreover, we determined the composition of intra- and extracellular dissolved free amino acids (DFAA) in all three species using high performance liquid chromatography (HPLC).

Our results indicate that marine AOA exude a suite of organic compounds with potentially varying reactivity, dominated by nitrogen-containing compounds. A significant fraction of the released DOM consists of ecologically important labile compounds, which are typically limiting in open ocean waters, including amino acids, thymidine and vitamins (B2 and B5). Particularly in the aphotic layers of the ocean where phytoplankton are essentially absent, “dark primary production” might play a crucial role in DOM formation and provide substrates for heterotrophic prokaryotes.

## Material and Methods

### Culture conditions

Axenic cultures of *Nitrosopumilus adriaticus* NF5^T^ (=JCM 32270^T^), *Nitrosopumilus piranensis* D3C^T^ (=JCM 32271^T^ =DSM 106147^T^) *and Nitrosopumilus maritimus* SCM1^T^ (=ATCC TSD-97^T^ =NCIMB 15022^T^) were grown in SCM medium as previously described^38^, with addition of 5 U mL^−1^ catalase. All plastic and glassware used were rinsed with acidified ultrapure water (Milli-Q, HCl analytical grade, pH 2), and glassware was combusted at 500°C for 5h. Cultures were grown in 2 L glass bottles for targeted and untargeted exometabolomics analysis and in 30 mL polypropylene tubes (Sterilin, Thermo Fisher Scientific) for dissolved free amino acid (DFAA) analysis. Growth was monitored by measuring nitrite production and cell abundance as previously described^38^.

### Untargeted and targeted exometabolomics

Culture supernatants of 1-2 L of culture were obtained via centrifugation (14,000x *g*, 10°C, for 1h) and subsequent filtration twice through 0.2 μm polycarbonate filters (GTTP, Millipore). DOM was extracted from culture supernatants by solid phase extraction (SPE) to remove salts (described in detail in ref.^39^). Briefly, supernatants were acidified to pH 2 (HCl, analytical grade) and extracted on PPL sorbent cartridges (1 g, Agilent, Waldbronn, Germany). Subsequently, cartridges were rinsed with acidified (pH 2) ultrapure water, air-dried, eluted with 6 mL methanol into amber glass vials and stored at −20°C. Procedural blanks were prepared by processing cell-free culture medium in the same way. Three biological replicates per archaeal strain were extracted. Due to the high culture volumes required, biological replicates were performed consecutively and culture medium blanks were generated for every medium batch (Supplementary Fig. 1). Dissolved organic carbon (DOC) concentrations of the solid-phase extracted DOM were quantified as described in ref^40^.

**Fig. 1.**
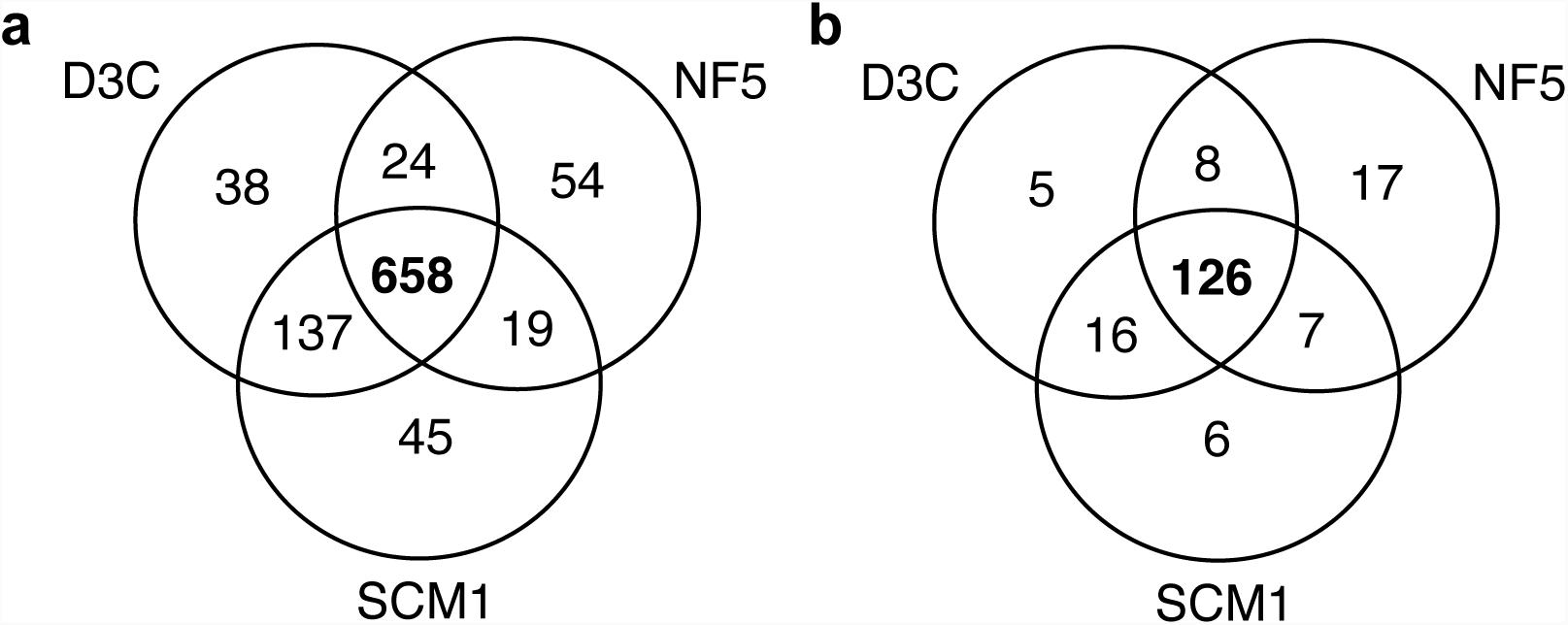
Venn diagram of detected molecular masses (**a**) and assigned formulae (**b**) present in *N. piranensis* D3C, *N. adriaticus* NF5, and *N. maritimus* SCM1.

Untargeted mass spectrometric analysis of DOM extracts was performed on a solariX Fourier transform ion cyclotron resonance mass spectrometer with a 15 Tesla magnet (Bruker, Daltonics, Bremen, Germany). The system was equipped with an electrospray ionization source (ESI, Bruker Apollo II, Daltonics, Bremen, Germany) applied in negative ionization mode. Prior to injection, methanol extracts were mixed with ultrapure water (50:50 v/v) and subsequently diluted with methanol: ultrapure water (50:50 v/v) according to the extracted volume (1:1 for 1 L, 1:2 for 2 L). For each measurement, 300 scans were accumulated in the mass window of 92 to 1,000 Da. Spectra were calibrated internally with a reference mass list using the Bruker Daltonic Data Analysis software Version 4.0 SP4. Instrument performance was verified using an in-house reference sample of North Equatorial Pacific Intermediate Water (NEqPIW) as previously described^40,41^. Technical triplicates were measured for every sample and only peaks detected in two out of three runs were kept in the dataset. Detected mass to charge (m/z) ratios were processed by applying a customized routine Matlab script and molecular formulae were assigned using the elements C, H, O, N, S, and P according to the guidelines established previously^42^. Masses detected in only one sample were removed as well as those with a maximum signal-to-noise ratio < 4 across all samples. Data were normalized to the sum of all peak intensities and only masses detected in two biological replicates were considered. Peaks of procedural blanks were removed when they were either present in more than half of all replicates (technical + biological) and/or if average peak intensities were higher than twice the average sample intensity. Assigned formulae were categorized into compound classes^28^ and carboxyl-rich alicyclic molecules (CRAMs) were identified as previously reported^43^. The exometabolome of the three archaeal species obtained by FT-ICR-MS was compared with the DOM from the water column of the North Atlantic acquired and analysed with the same approach^4^. The unfiltered and filtered datasets can be found in Supplementary data file 1.

For targeted analysis, DOM extracts were dried down and re-dissolved in 95:5 (v/v) water:acetonitrile and deuterated biotin was added as an injection standard (final concentration 0.05 mg mL^−1^). All three biological replicates from each *Nitrosopumilus* strain and two out of five media blanks were selected for targeted analysis. Samples were analysed by ultra-high performance liquid chromatography (Accela Open Autosampler and Accela 1250 Pump, Thermo Scientific) coupled to a heated electrospray ionization source (H-ESI) and a triple quadrupole mass spectrometer (TSQ Vantage, Thermo Scientific) operated under selected reaction monitoring (SRM) mode. Chromatographic separation was performed on a Waters Acquity HSS T3 column (2.1 × 100 mm, 1.8 μm) equipped with a Vanguard pre-column and maintained at 40°C. The column was eluted with (A) 0.1% formic acid in water and (B) 0.1% formic acid in acetonitrile at a flow rate of 0.5 mL min^−1^ as previously described^44^. Separate autosampler injections of 5 μL each were made for positive and negative ion modes. The samples were analysed in a random order with a pooled sample run after every six samples. The mass spectrometer was operated in selected reaction monitoring (SRM) mode. Optimal SRM parameters (s-lens, collision energy) for each target compound were optimized individually using an authentic standard as previously described^44^. Two SRM transitions per compound were monitored for quantification and confirmation. Eight-point external calibration curves based on peak area were generated for each compound. The resulting data were converted to mzML files using the msConvert tool^45^ and processed with MAVEN^46^. Compounds were considered when they were present in at least two out of three biological replicates and absent in media blanks.

### Determination of dissolved free amino acids (DFAA)

For the analysis of extracellular DFAA, 1 mL of culture was filtered through 0.2 μm syringe filters with PVDF membrane (Whatman^®^ Puradisc 13) and the filtrate was stored in 1.5 mL combusted amber glass vials at −20°C until analysis. For the analysis of intracellular DFAA, cells were harvested via centrifugation as described above and 1.5 mL Milli-Q was added to the cell pellets. Cells were subsequently lysed with 4 freeze/thaw cycles and the lysate was filtered through 0.1 μm syringe filters (Millipore Durapore, 25 mm). Concentrations of extracellular DFAA were determined throughout the growth of the three strains and intracellular DFAA were determined at late exponential growth phase in all three strains.

Intra- and extracellular DFAA were analysed using a high performance liquid chromatography (HPLC) system (Agilent 1260 Infinity Bioinert) equipped with a Zorbax Sq-Aq Analytical guard column (4.6 × 12.5 mm, 5 μM) and Zorbax Eclipse AAA Rapid resolution column (4.6 × 150 mm, 3.5 μM; Agilent Technologies, Santa Clara, USA). To 1 mL of sample, 75 μL borate buffer (0.4 M, pH 10.2) and 5 μL OPA reagent (5061-3335, Agilent Technologies) were added for the derivatization reaction (27°C, 2 min). The injection volume was 100 μL and separation was achieved by applying a flow rate of 0.8 mL min^−1^ under a constant column temperature of 25°C using a modified gradient to reduce interference of high ammonium concentrations (Supplementary table 1) (Taubner et *al., unpublished*). Excitation and emission wavelengths of the fluorescence detector were 340 nm and 450 nm, respectively, with gain factor 12 for quantification. For peak identification and area calculation, a primary amino acid standard (A2161, Sigma-Aldrich) was prepared in the range of 1 nM to 1 μM. Spectra were analysed with the software Agilent ChemStation (Agilent Technologies).

**Table 1.**
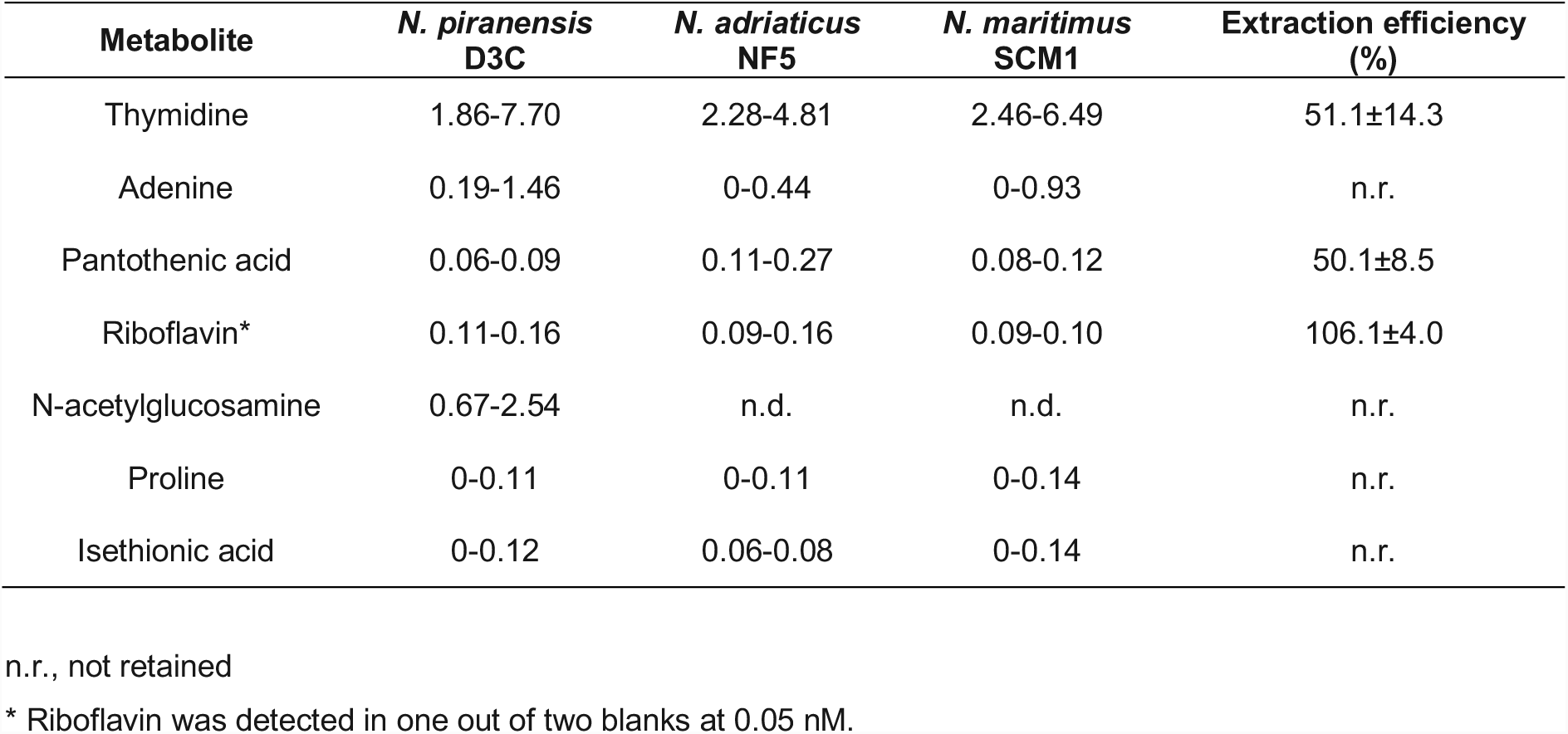
Targeted exometabolomics analysis of SPE-extracted DOM from *N. piranensis* D3C, *N. adriaticus* NF5 and *N. maritimus* SCM1. Metabolite concentrations are given in nM and represent the minimum and maximum values detected in three biological replicates per strain. Extraction efficiencies of metabolites in Milli-Q water can be found in ref.^80^ and the complete list of metabolites has previously been reported in ref.^44^.

| Metabolite | *N. piranensis*<br>D3C | *N. adriaticus*<br>NF5 | *N. maritimus*<br>SCM1 | Extraction efficiency<br>(%) |
| --- | --- | --- | --- | --- |
| Thymidine | 1.86-7.70 | 2.28-4.81 | 2.46-6.49 | 51.1±14.3 |
| Adenine | 0.19-1.46 | 0-0.44 | 0-0.93 | n.r. |
| Pantothenic acid | 0.06-0.09 | 0.11-0.27 | 0.08-0.12 | 50.1±8.5 |
| Riboflavin* | 0.11-0.16 | 0.09-0.16 | 0.09-0.10 | 106.1±4.0 |
| N-acetylglucosamine | 0.67-2.54 | n.d. | n.d. | n.r. |
| Proline | 0-0.11 | 0-0.11 | 0-0.14 | n.r. |
| Isethionic acid | 0-0.12 | 0.06-0.08 | 0-0.14 | n.r. |

n.r., not retained
\* Riboflavin was detected in one out of two blanks at 0.05 nM.

Limits of detection (LOD) and quantification (LOQ), as well as the recovery of individual amino acids in Milli-Q water and culture medium can be found in the Supplementary information. Extracellular methionine and valine concentrations could not be reliably quantified until late exponential/early stationary growth phase of the cultures due to the interference with ammonium in the culture medium. Tyrosine could never be reliably quantified in extracellular samples due to ammonium interference and the presence of three peaks with similar retention times. Lysine could only be measured in the culture supernatant of *N. adriaticus* NF5, due to other compound(s) with similar retention times present at micromolar concentrations in the medium of *N. piranensis* D3C and *N. maritimus* SCM1. Aspartic acid, serine, glutamine and histidine often appeared as small double peaks, which were integrated together for consistency.

## Results and Discussion

### Composition of solid phase extracted-DOM produced by ammonia-oxidizing archaea

Dissolved organic matter (DOM) was extracted from culture supernatant of three strains of ammonia-oxidizing archaea (AOA), *Nitrosopumilus adriaticus* NF5, *Nitrosopumilus piranensis* D3C and *Nitrosopumilus maritimus* SCM1. Dissolved organic carbon (DOC) concentrations of solid-phase extracted DOM (SPE-DOM) from archaeal cultures harvested during late exponential growth ranged from 3.5 to 5.0 μM, which was on average 2-3 times higher than DOC concentrations of medium blanks (data not shown). We were not able to determine the extraction efficiencies due to the high concentrations of organic HEPES buffer (10 mM) in the culture medium, which was not retained by the PPL cartridges. Extraction efficiencies of DOC from seawater are ∼60% on average, typically exhibiting a C/N ratio similar to the original DOM^47^. However, DOC extraction efficiencies in cultures of heterotrophic bacteria and phytoplankton are typically lower, ranging from 2 to 50%^26,27,48^. The amount of SPE-DOC released by the three archaeal strains was significantly lower than has been previously determined for heterotrophic bacteria (3.5-5.0 μM compared to 144-511 μM as in ref. ^28^). Given the energetic differences between the oxidation of ammonia (ΔG=-271 kJ mol^−1^ NH_3_) and glucose (−2883 kJ mol^−1^ glucose), it is not surprising that these organisms are not releasing as much (fixed) carbon as compared to heterotrophic bacteria. The carbon yield from nitrification (fmol C fixed cell^−1^ d^−1^/ fmol N oxidized cell^−1^ d^−1^) of *N. adriaticus* is ∼0.1, amounting to 10 moles of NH_3_ which need to be oxidized for every mole of carbon fixed (Bayer *et al*., *submitted*). Consequently, for autotrophic ammonia-oxidizers, it is much more costly to produce these extracellular compounds than for heterotrophic bacteria. The fraction of DOC exuded by *Nitrosopumilus* spp. was estimated to account for ∼4 - 50% of the fixed carbon in this study. The large variation results from uncertainties regarding the extraction efficiency required to remove the salts.

A total of 5442 resolved masses of singly charged, intact compounds were detected with FT-ICR-MS. After stringent filtering of masses present in medium controls, 976 masses remained that were present in at least two technical and biological replicates per strain (see Material and Methods for details) (Fig. 1a). Thereof, 185 masses could be assigned to molecular formulae of which 126 were shared by all three strains (Fig. 1b). Of these shared exometabolites, 78.6% contained at least one nitrogen atom, which was much higher than the average percentage of nitrogen-containing compounds in DOM extracted from the North Atlantic (Fig. 2a,c). Previous studies showed that DOM released by heterotrophic bacteria is characterized by a high abundance of molecules with heteroatoms (P, N, S)^28,48^. In comparison to the exometabolome of the marine bacterium *Pseudovibrio* sp. FO-BEG, however, S- and P-containing compounds were much less abundant in the exometabolomes of *Nitrosopumilus* spp.^28^, potentially reflecting the dominance of nitrogen metabolism in AOA (Fig. 2b,c).

**Fig. 2.**
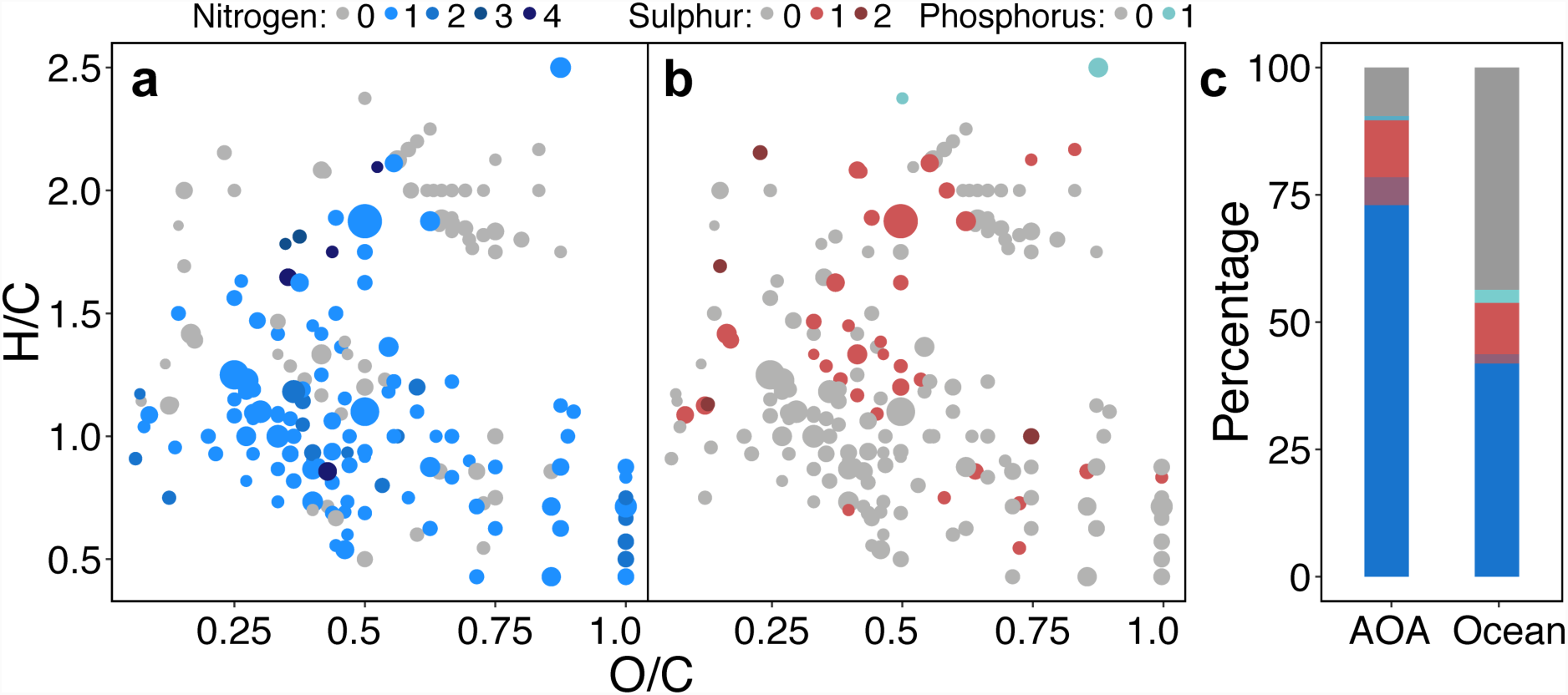
Van Krevelen plots of masses with assigned formulae derived from *Nitrosopumilus* spp. exometabolomes containing nitrogen (**a**), sulphur and phosphorus (**b**) heteroatoms. Dot size corresponds to average relative peak intensity. (**c**) Percentage of assigned formulae in ocean DOM and archaeal DOM containing nitrogen, sulphur or phosphorus atoms. The presence of two different heteroatoms in one molecular formula is indicated by the overlay of both colours.

The number of detected masses with assigned molecular formulae was much lower in SPE-DOM of *Nitrosopumilus* spp. than observed for heterotrophic bacteria using comparable methodology^27,28^. However, it remains unclear whether the exometabolome of chemolithoautotrophic *Nitrosopumilus* spp. is generally less diverse as compared to that of heterotrophic bacteria, or whether *Nitrosopumilus* spp. produces fewer compounds that are retained by SPE columns. SPE-DOM is primarily concentrated based on polarity when using PPL cartridges^39^, and is relatively rich in low-molecular weight compounds and carboxyl-rich alicyclic molecules (CRAM). CRAM accounted for 31% of all archaeal DOM molecules (Fig. 3a), which is comparable to previously reported CRAM production by bacterioplankton^48^. It has been suggested that CRAM are largely comprised of decomposition products of biomolecules, representing a major refractory component of oceanic DOM^43^. However, the rapid generation of CRAM by archaeal and bacterial metabolism (ref.^48^, this study) challenges their potentially refractory nature. If such a large proportion of freshly produced DOM was refractory, then the oceans would be filled almost exclusively by CRAM. Furthermore, recent evidence suggests that the low concentrations of individual compounds rather than their inherent recalcitrance prevents prokaryotic consumption of a substantial fraction of DOM in the ocean^49^. SPE-DOM from *Nitrosopumilus* spp. supported the growth of a heterotrophic alphaproteobacterium (Supplementary Fig. 2), suggesting that at least a fraction of the released DOM retained on SPE columns might be used as substrate by heterotrophic bacteria.

**Fig. 3.**
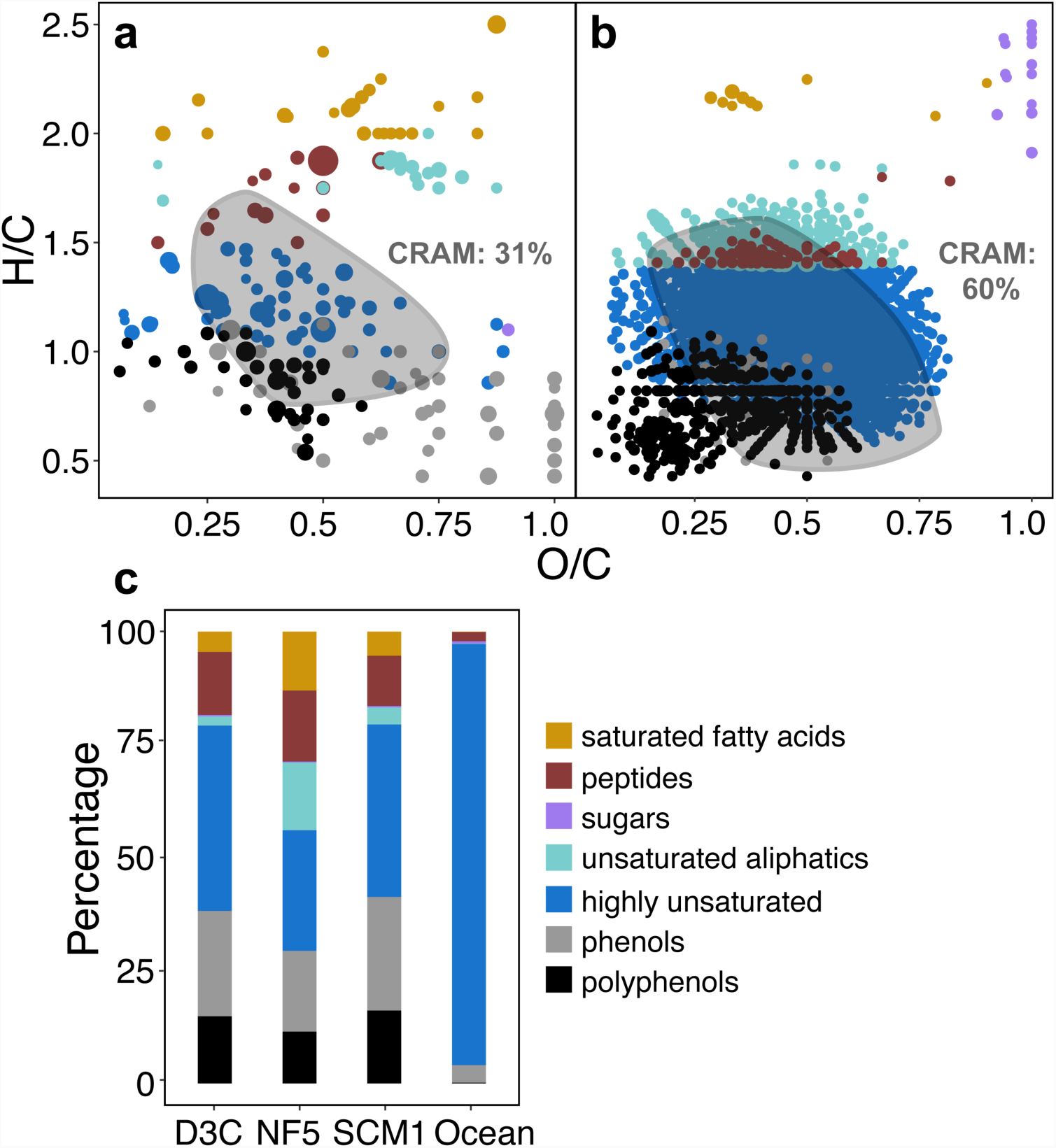
Categorization of assigned formulae obtained from *Nitrosopumilus* spp. DOM (**a**) and ocean DOM (**b**) into compound classes (for definition of categories see Material and Methods). Dot sizes correspond to average relative peak intensities and range of O/C and H/C ratios corresponding to carboxyl-rich alicyclic molecules (CRAM) is indicated by grey circles. (**c**) Comparison of the DOM compositions of three *Nitrosopumilus* strains (*N. piranensis* D3C, *N. adriaticus* NF5 and *N. maritimus* SCM1) with ocean DOM signatures normalized to relative peak intensities.

While oceanic DOM is dominated by highly unsaturated compounds, *Nitrosopumilus*-derived DOM contained a high proportion of compounds with molecular formulae characteristic for phenols, polyphenols, highly unsaturated compounds, unsaturated aliphatics, peptides and saturated fatty acids (Fig. 3a,c). The exometabolomes of AOA contained relatively fewer sugars and peptides and a higher percentage of phenols compared to *Pseudovibrio* sp. FO-BEG^28^. The production of phenols and polyphenols has been associated with antioxidant activity, suggested to be caused by increased oxidative stress^28^. AOA were shown to produce high amounts of the oxidant H_2_O_2_ as a result of their metabolic activity, which ultimately inhibits their growth^25^. Although, the H_2_O_2_-detoxifying enzyme catalase was initially added to the culture medium, catalase activity gradually decreases with time, potentially increasing the exposure to oxidative stress in *Nitrosopumilus* spp. cultures. A relatively high proportion of the assigned molecular formulae of the exometabolomes of all three strains was assigned to saturated fatty acids (Fig. 3a,c). In contrast to Bacteria, archaeal membrane lipids consist of isoprenoids and do not contain fatty acids^50^. However, fatty acids have been detected in several Crenarchaeota species and in halorhodopsins of *Haloarchaea*^51,52^. Recently, it was suggested that Archaea potentially synthesize fatty acids by reversing the β-oxidation pathway^53^. *Nitrosopumilus* spp. genomes encode putative enzymes involved in this proposed pathway, including enoyl-CoA hydratase, 3-hydroxyacyl-CoA dehydrogenase and acetyl-CoA C-acetyltransferases, as well as for a distant homolog of FadE that could potentially exhibit acyl-CoA dehydrogenase activity^53^.

### Release of ecologically relevant metabolites by three *Nitrosopumilus* strains

Targeted exometabolomics revealed that only a small fraction of the 92 specific compounds tested was identified in SPE-DOM of *Nitrosopumilus* spp. (Table 1). A major feature observed in the targeted analysis of the extracellular metabolite fractions was the presence of the nucleoside thymidine, which was the most abundant identified metabolite in all biological replicates of each strain, ranging between 3.72 and 15.40 nM when corrected for extraction efficiency (Table 1). The amount of thymidine in SPE-DOM would represent ∼10% of the total cellular thymidine content contained in DNA in the cultures (calculated based on the dsDNA weight of 1 fg Mb^−1^ and a G+C content of 34 mol%). However, cultures were harvested in late exponential growth phase and no decrease in cell abundance was observed (data not shown), suggesting that thymidine was released by growing cells. Thymidine has also been reported as a dominant molecule in the exometabolome of the cyanobacterium *Synechococcus elongatus*^54^. Like Synechococcus, the genomes of members of the *Nitrosopumilus* genus lack thymidine kinase, which phosphorylates thymidine to thymidine monophosphate (TMP) in the thymidine salvage pathway. Instead, uridine 5’-monophosphate (UMP) is converted to TMP by thymidylate synthase for DNA synthesis. Thus, thymidine released by TMP hydrolysis or DNA degradation cannot be metabolized and is likely excreted as waste product^54^. Thymidine represents a valuable substrate for heterotrophic bacterioplankton and is readily taken up and incorporated into DNA, a feature that has been applied to measure heterotrophic prokaryotic production in aquatic environments^55,56^.

Minor components of the exometabolomes might be ecologically important even at low concentrations, such as pantothenic acid (vitamin B5) and riboflavin (vitamin B2), which were found at concentrations of 120-540 pM and 90-160 pM (corrected for extraction efficiencies), respectively, in all three cultures. B vitamins are some of the most commonly required biochemical cofactors for cellular metabolism, yet their concentrations vary from undetectable to a few picomoles per liter in ocean waters^57,58^. Adenine, proline and isethionic acid were detected in the exometabolomes of all three strains, yet not in every biological replicate (Table 1). These compounds are typically not retained by SPE but probably present at high amounts in our study, rendering their reliable quantification difficult. Furthermore, the amino sugar *N*-acetylglucosamine was only detected in SPE-DOM of *N. piranensis*, yet in all three biological replicates (Table 1), indicating that *N. piranensis* D3C produced higher amounts of this metabolite as compared to the other two species. While Archaea lack peptidoglycan, which largely consists of *N*-acetylglucosamine and *N*-acetylmuramic acid, archaeal S-layers are often heavily glycosylated and glycan composition has been shown to vary even between very closely related archaeal species^59^.

We also determined concentrations of dissolved free amino acids (DFAA) in the culture medium throughout the growth of all three *Nitrosopumilus* strains. Total DFAA concentrations ranged between 261-454 nM at the end of exponential growth and net cell-specific DFAA release rates were between 2.6 and 16.4 fmol cell^−1^ d^−1^ (with *N. piranensis* exhibiting the highest rates) (Table 2), which is comparable to rates previously reported for marine phytoplankton species (0.17-38 fmol cell^−1^ d^−1^)^60,61^. Surprisingly, *Nitrosopumilus* spp. Released DFAA at higher rates than two heterotrophic alphaproteobacterial strains of the marine *Roseobacter* clade (concentrations were below the detection limit of 0.5 nM)^27^. Furthermore, the extracellular DFAA composition appeared to be somewhat connected to the phylogenetic relatedness of the three species, as shown for different phytoplankton species^26,62^. *N. adriaticus* NF5 released higher concentrations of serine, whereas *N. piranensis* D3C and *N. maritimus* SCM1 released higher concentrations of methionine and asparagine (Fig. 4a, Supplementary table 3). When exposed to oxidative stress, cell-specific DFAA release rates increased, in particular those of glutamine and threonine, as compared to cultures grown under non-stress conditions (Supplementary Fig. 3).

**Table 2.** Extracellular total dissolved free amino acid (DFAA) concentrations produced by the three *Nitrosopumilus* strains (*N. piranensis* D3C, *N. adriaticus* NF5 and *N. maritimus* SCM1). Standard deviations of triplicate measurements are shown in parentheses.

| Strain | Time<br>(d) | Cells<br>($\times 10^6$ mL<sup>-1</sup>) | Total DFAA<br>(nM) | +Val, Met<br>(nM) | Release rates<sup>‡</sup><br>(fmol cell<sup>-1</sup> d<sup>-1</sup>) |
| --- | --- | --- | --- | --- | --- |
| NF5 | 0 | 3.1 |  |  |  |
| NF5 | 3 | 10.7 | 26 (17) |  | 3.4 (2.2) |
| NF5 | 5 | 23.5 | 135 (24) |  | 8.5 (0.5) |
| NF5 | 7 | 31.1 | 164 (33) | 209 (36) | 3.8 (1.2) |
| NF5 | 9 | 34.6 | 214 (44) | 261 (40) | 14.3 (1.1) |
| D3C | 0 | 2.1 |  |  |  |
| D3C | 3 | 5 | 44 (42) |  | 15.2 (14.5) |
| D3C | 6 | 11.7 | 154 (27) |  | 16.4 (2.2) |
| D3C | 9 | 29.4 | 346 (51) | 454 (84) | 9.0 (4.5) |
| SCM1 | 0 | 1.8 |  |  |  |
| SCM1 | 3 | 7.2 | 14 (14) |  | 2.6 (2.6) |
| SCM1 | 6 | 29.4 | 164 (63) | 239 (73) | 6.8 (2.2) |
| SCM1 | 8 | 48.2 | 235 (22) | 320 (34) | 3.8 (2.2) |

<sup>‡</sup> Valine and methionine were excluded from per-cell DFAA release rates as they could only be quantified in the absence of NH<sub>4</sub><sup>+</sup>/NH<sub>3</sub>, corresponding to the end of exponential growth (see Material and Methods section)

**Fig. 4.**
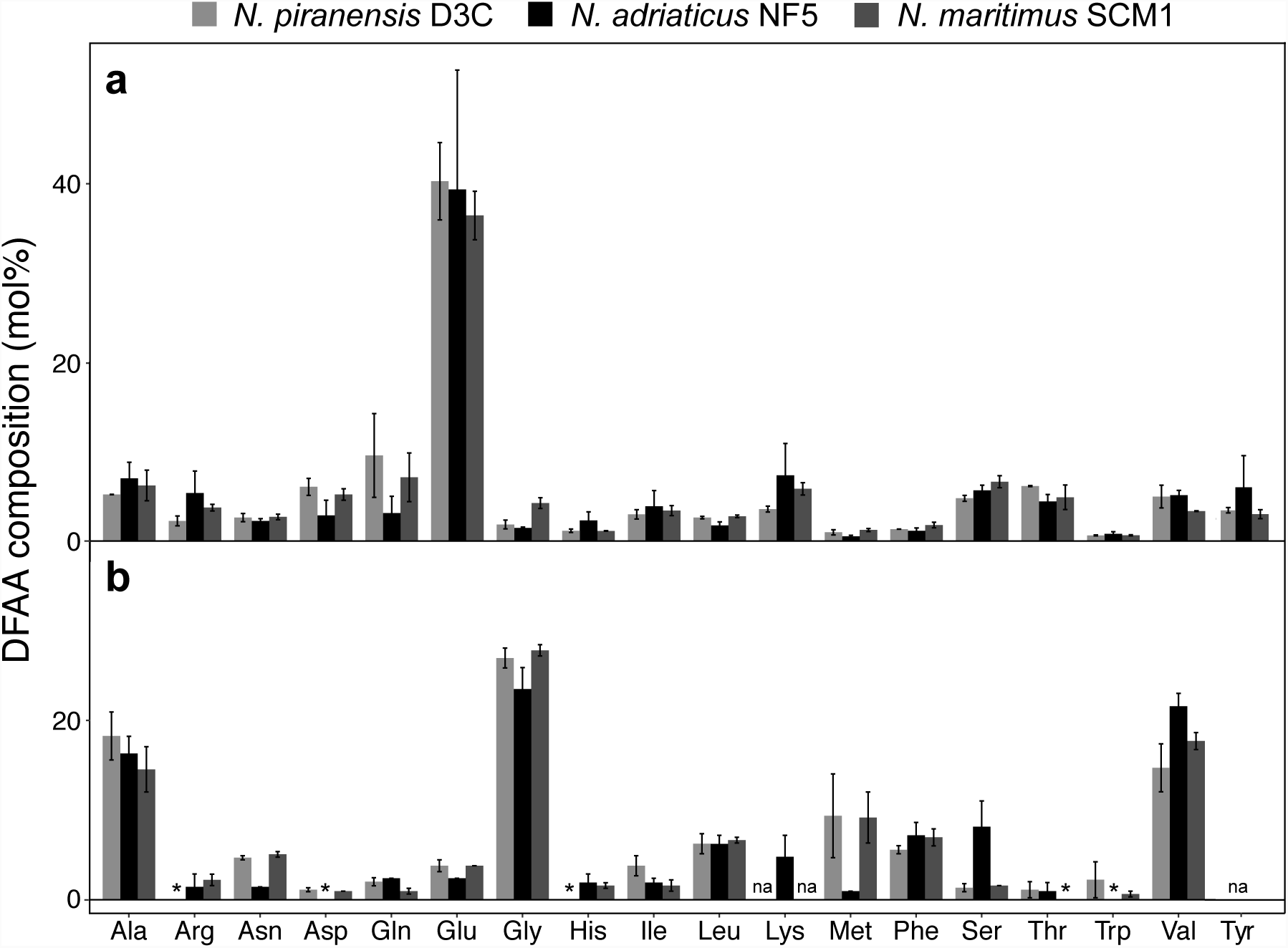
Intracellular (**a**) and extracellular (**b**) dissolved free amino acid (DFAA) composition of *N. piranensis* D3C, *N. adriaticus* NF5, and *N. maritimus* SCM1. Error bars represent standard deviations of measurements from triplicate cultures. *, values below detection limit; na, not available.

### Potential mechanisms of DFAA release by ammonia-oxidizing archaea

The fraction of DOM released by phytoplankton is highly variable accounting for up to 70% of the total photosynthetically fixed carbon^6,18^. Phytoplankton exudation has largely been interpreted as the active release of excess photosynthates when carbon fixation exceeds incorporation into cell material^20^. However, Bjørnson speculated already more than 30 years ago that diffusion rather than overflow metabolism might be the responsible mechanism of the photosynthetic extracellular release^63^.

Among the three *Nitrosopumilus* strains, the intracellular DFAA composition was highly similar, with glutamic acid dominating the intracellular DFAA pool (35-50 mol%) (Fig. 4a). With the exception of glutamic acid, the intracellular DFAA composition largely reflected the average amino acid composition encoded by *Nitrosopumilus* spp. genomes (Supplementary Fig. 4). Ammonia is typically incorporated into biomolecules through glutamate and thus, glutamate/glutamic acid is present at elevated concentrations in most cells. Intriguingly, glutamic acid was largely absent from the extracellular DFAA pool of growing cells (Fig. 4b). Instead, hydrophobic amino acids, including alanine, glycine, valine, leucine, isoleucine, phenylalanine, methionine and tryptophan made up ∼80% of the extracellular DFAA of *Nitrosopumilus* spp. during late exponential growth phase. Additionally, proline, which is also highly hydrophobic, was detected via targeted exometabolomics (Table 1). The permeability of the lipid bilayer for hydrophobic amino acids might be up to 100 times higher as compared to hydrophilic amino acids, as indicated in experiments with artificial lipid vesicles^64^. Furthermore, amino acid transport mutants of the cyanobacterium *Anabaena* were shown to release a mixture of hydrophobic amino acids, supporting the hypothesis of passive diffusion^65^. We did not identify putative amino acid transporters in the proteomes of either species (Bayer et *al., unpublished*), suggesting that the extracellular DFAA composition is determined by selective release rather than selective re-uptake of amino acids by the cells.

The release of most amino acids followed the growth patterns of each strain, as indicated by nitrite production (Fig. 5, Supplementary Fig. 4), with the exception of glycine which was continuously released at high rates. The cleavage of L-serine to glycine results in the formation of 5,10-methylenetetrahydrofolate (from tetrahydrofolate), which represents an important one-carbon unit donor for pyrimidine, purine and methionine biosynthesis^66^. The decoupling of glycine release (and its precedent production) from ammonia oxidation might indicate that pyrimidine/purine biosynthesis is maintained at a similar level by the cell even when reaching stationary phase.

**Fig. 5.**
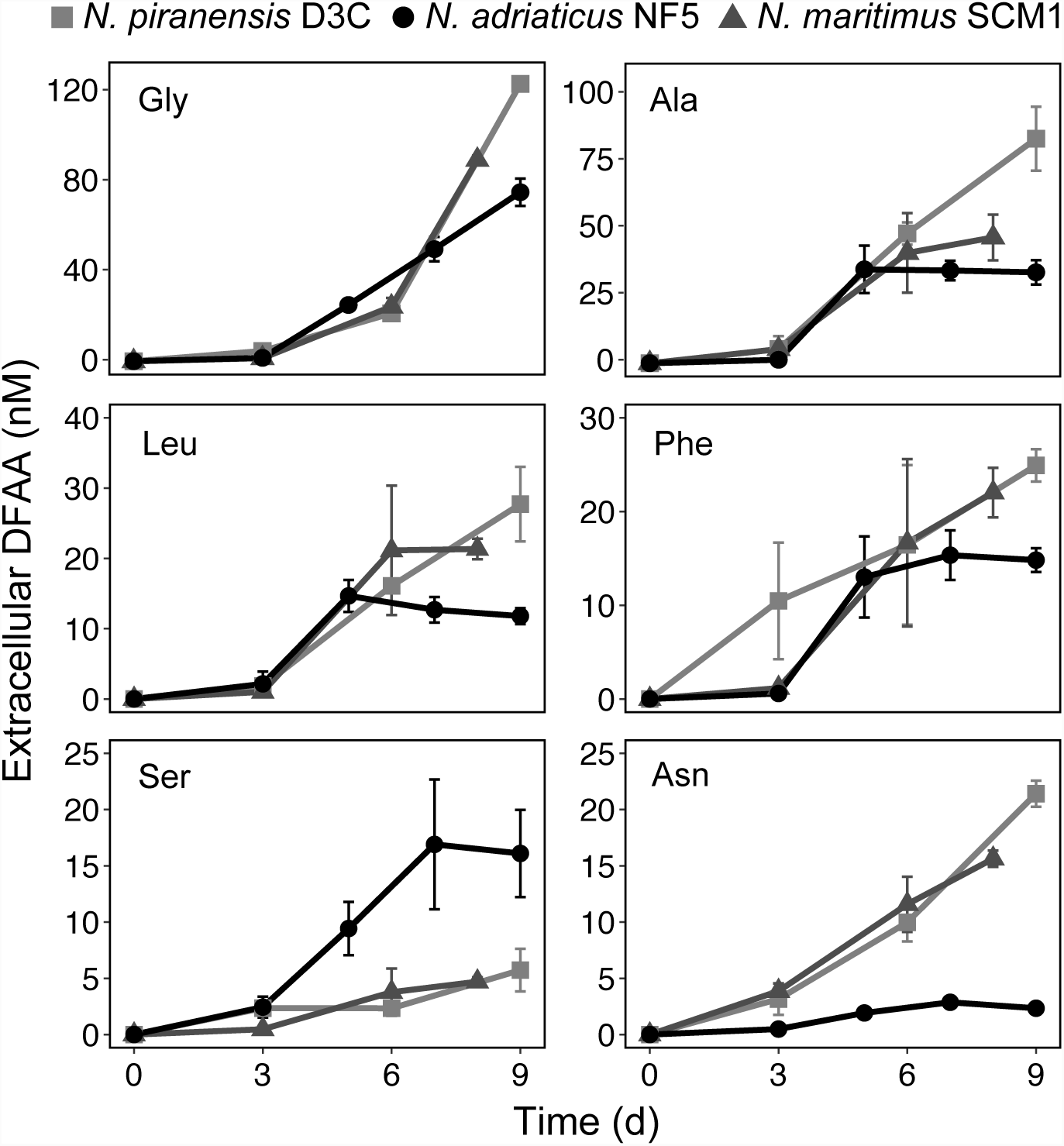
Net release of dissolved free amino acids (DFAA) throughout the growth of three *Nitrosopumilus* strains. T0 concentrations were subtracted from all time points. Note that the scale bars are different between different amino acid species. Error bars represent standard deviations of measurements from triplicate cultures.

### DOM release by ammonia-oxidizing archaea: Implications for the microbial food web

Phytoplankton are considered the major producers of DOM in the surface ocean, accounting for a large fraction of labile carbon flux which fuels secondary production^20,21^. Below the sunlit surface layers of the ocean, chemolithoautotrophy is a widespread strategy and this “dark primary production” has a major impact on global carbon cycling^67^. Our data suggests that AOA might release a substantial fraction of the fixed carbon to the ambient environment in the form of amino acids and other metabolites, which has not been taken into account thus far. Quantifying the chemolithoautotrophic contribution to the DOM pool and its turnover represents a potentially important aspect to improve models of the global ocean carbon cycle.

Furthermore, the inability of an organism to synthesize a particular compound required for its growth (auxotrophy) appears to be more widespread than previously assumed^68^. Exploiting a resource produced by another organism rather than *de novo* synthesizing a specific compound has been suggested to provide a selective advantage leading to reductive genome evolution (“Black Queen Hypothesis”)^69^. The auxotrophy for B vitamins in phytoplankton has been known for decades^70^. However, a recent survey of bacterial genomes (from various environments) revealed that only 1% encoded the complete pathways for all 20 essential amino acids^71^.

The heterotrophic marine bacterium *Pelagibacter ubique*, a member of the SAR11 clade, known as the most abundant organisms on the planet^72^, lacks biosynthetic pathways for various vitamins (B_1_, B_5_, B_6_, B_7_, B_12_) and requires pyruvate, glycine and organic sulphur (e.g., methionine) for growth^73–75^. Recently, it was shown in co-culture experiments that photoautotrophic *Prochlorococcus* strains can fulfill some of the unique metabolic requirements of SAR11^76^. Additionally, members of the *Roseobacter* group have been suggested as important suppliers of so-called “public goods” by releasing growth factors as well as biosynthetic precursors, including various vitamin B precursors^27^. B vitamins are limiting microbial production in large areas of the ocean, thus potentially constraining rates of primary production and carbon sequestration^58^. AOA are prototrophic for many B vitamins and have previously been suggested to play a critical role in the microbial metabolic activity in meso- and bathypelagic waters by supplying these essential nutrients^77,78^. Our results provide the first evidence for the release of B vitamins by marine AOA. Curiously, concentrations of multiple B vitamins were shown to be enriched primarily in the upper mesopelagic zone^58^ where AOA typically exhibit peaks of abundance and activity^79^.

## Conclusions

This study characterizes the exometabolome of autotrophic ammonia-oxidizing archaea (AOA). Untargeted and targeted exometabolomic profiling of DOM from three model strains of AOA suggests that these abundant chemolithoautotrophs might influence the chemical composition of the oceanic DOM. Our results provide evidence that AOA release a suite of organic compounds with potentially varying reactivity. However, the rapid generation of carboxyl-rich alicyclic molecules (CRAM) by AOA challenges the paradigm that CRAM are solely representative for old and refractory compounds in the global ocean.

The release of fixed carbon represents an unprecedented link between chemolithoautotrophic production and heterotrophic consumption of DOM in the ocean and has important implications for the role of AOA in the microbial carbon pump. Furthermore, the release of ecologically and biochemically relevant metabolites could potentially be crucial for microbes that are auxotrophic for some of these compounds, including members of the globally abundant and ubiquitous SAR11 clade. Future efforts in quantifying the chemolithoautotrophic contribution to the marine DOM pool and its turnover are needed in order to improve models of the global carbon cycle, ultimately contributing to our understanding of the factors influencing the long-term storage of carbon in the ocean’s interior.

## Acknowledgements

We thank Ina Ulber for TOC measurements, Katrin Klaproth for FT-ICR-MS data processing and Christian Baranyi for technical assistance with cultivations. We also thank Elisabeth Clifford and Barbara Mähnert for establishing a HPLC method that is less affected by high ammonium concentrations. We are grateful to the FT-MS facility at Woods Hole Oceanographic Institute (WHOI) for targeted metabolomics analysis and Krista Longnecker for helpful comments. BB was supported by the Uni:docs Fellowship of the University of Vienna, and the Austrian Science Fund (FWF) DK+ project “Microbial Nitrogen Cycling” (W1257-B20) and the FWF project ARTEMIS (P28781-B21), both to GJH. This work is part of the requirements for the fulfilment of the PhD degree of BB.

## Author contributions

BB, RLH and GJH designed the research. BB performed all laboratory experiments and measured samples on the HPLC and FT-ICR-MS. MJB contributed to amino acid sampling and analysed amino acid chromatograms. BN-O, JN and TD contributed to FT-ICR-MS data acquisition and analyses. Targeted metabolomics measurements were performed by the Woodshole Oceanographic Institute. The manuscript was written by BB, with input from all co-authors.

