## Supplementary Information for "Ammonia-oxidizing archaea release a suite of organic compounds potentially fueling prokaryotic heterotrophy in the ocean"

### Supplementary Information (SI)

#### Supplementary Methods

*Accuracy and precision of dissolved free amino acid (DFAA) determination with HPLC*

Spent culture medium (containing µM ammonium concentrations) and Milli-Q test samples were amended with three concentrations of amino acid standard mix (5 nM, 50 nM, 500 nM). Five replicates for each concentration were measured. Recovery of DFAA in Milli-Q and culture medium was determined by calculating the average recovery from amino acid standards (Table S2).

The recovery R (in %) was estimated as:

$$R = \frac{(\mathrm{Ca} - \mathrm{Cu})}{\mathrm{Cs} \times 100}$$

where Ca and Cu are the concentrations of the amino acids in amended and unamended samples, respectively, and Cs is the concentration of the amino acid standard added to the test samples.

*R* ranged from 92.1% to 115.4% in Milli-Q and from 59.5 % to 116.5 % in culture medium. *R* was generally more variable in culture medium and lower than in Milli-Q for most amino acids, except for asparagine, aspartic acid, glutamine, histidine, glycine, arginine and methionine (Table S2). The precision of the method was determined by calculating the relative standard deviation (RSD %) for the replicates. Limits of detection (LOD) and quantification (LOQ) were determined from five replicates and defined as signal-to-noise ratios of ≥ 3 and ≥10, respectively. Replicates were measured in different runs in order to capture the variability observed between runs. LOD and LOQ could only be determined in Milli-Q due to the amino acid background of spent culture medium (Table S2). The high LOD and LOQ of alanine are likely due to variable levels of alanine contamination in Milli-Q and/or from the air. Lysine could not be reliably quantified in most runs.

*Growth of the alphaproteobacterium* O. alexandrii *on SPE-DOM from* Nitrosopumilus *spp.*

*Oceanicaulis alexandrii* was grown in SCM medium (described in ref.[1]) amended with 100 µM amino acids (alanine, aspartic acid, glutamic acid, serine) in the dark at 30°C without shaking and growth was monitored by flow cytometry[2]. After reaching late exponential growth, *O. alexandrii* was starved for 3 d and then transferred into nine 20 mL flasks containing fresh SCM medium (without substrate addition). To three replicates, *Nitrosopumilus*-derived SPE-DOM and SPE-DOM from culture medium controls was added, respectively, and three replicates served as no substrate control incubations. The methanol extracts (1.2 mL each) were dried in a SpeedVac (Eppendorf, Concentrator Plus), subsequently re-dissolved in SCM medium and sterile-filtered through 0.1 µm syringe filters. DOM extracts were added to 20 mL of *O. alexandrii* culture each, resulting in a 20X concentration as compared to the initial DOC concentrations retained on SPE columns (ca. 40 µM of *Nitrosopumilus*-derived DOM and 20 µM of medium blank SPE-DOM).

#### Supplementary Results and Discussion

*DFAA release by three* Nitrosopumilus *species under $H_2O_2$ stress*

Axenic cultures of *Nitrosopumilus adriaticus* NF5, *Nitrosopumilus piranensis* D3C and *Nitrosopumilus maritimus* SCM1 were grown in SCM medium without addition of purified catalase which resulted in growth retardation and inhibition due to accumulation of $H_2O_2$ (Bayer *et al., unpublished*). In cultures exposed to $H_2O_2$ stress, the extracellular DFAA composition changed, resulting in an absolute increase in threonine and glutamine concentrations in the culture media of all three strains, as well as in a relative increase of histidine in *N. adriaticus* NF5 and *N. maritimus* SCM1 (Figure S3, Table S3). Exposure to $H_2O_2$ stress was accompanied by a stagnation in growth as previously reported (Bayer et *al.*, *submitted*). However, cell abundances did not decrease over the time course of the experiment, suggesting that increased release is not associated with cell death and biomass decay (Table S3). The increased release of DFAA could be due to membrane re-arrangement as a result of oxidative stress as described elsewhere (Bayer et *al.*, *submitted*), potentially leading to increased leakage of cytosolic metabolites. Alternatively, amino acids could be derived directly from the re-arranged membrane proteins since polar amino acids, generally found on the surface of a protein, were increasingly released under oxidative stress. However, the potentially non-efficient regulation of substrate uptake during stress conditions could also result in the secretion of non-metabolizable intermediates including amino acids.

#### Supplementary Figures and Tables

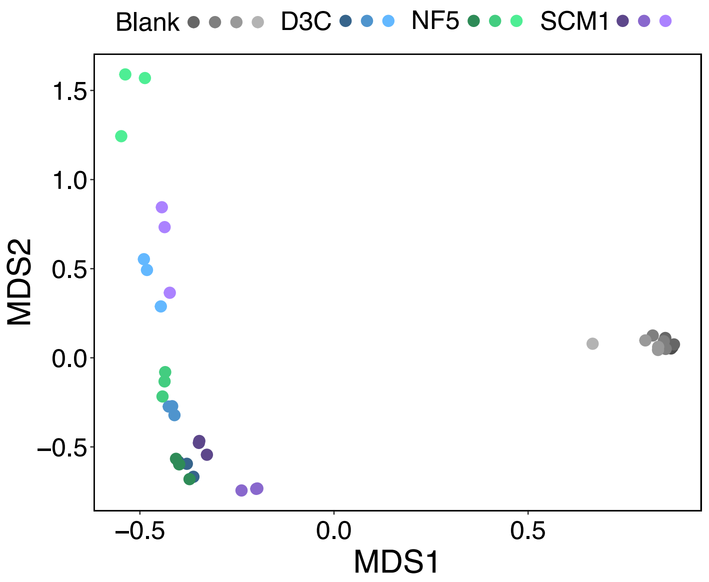


**Figure S1** Principal coordinates analysis (PCoA, = Multidimensional scaling, MDS) of all resolved masses of compounds detected in *Nitrosopumilus* strains (*N. piranensis* D3C, *N. adriaticus* NF5 and *N. maritimus* SCM1), as well as in all medium blanks. Bray-Curtis distance matrices were calculated using the normalized peak intensities and analysis was performed using the vegan package[3] in R[4]. Biological replicates are represented by different color shadings and technical replicates are represented by identical colors.

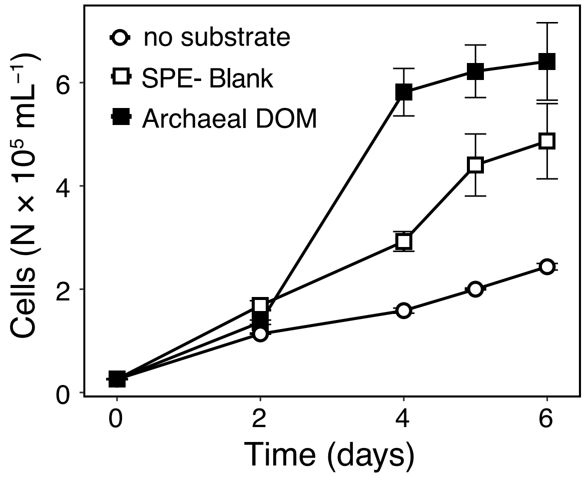


**Figure S2** Growth of the heterotrophic alphaproteobacterium *O. alexandrii* on SPE-DOM from archaeal culture supernatants and culture medium in comparison to control incubations without addition of organic carbon.

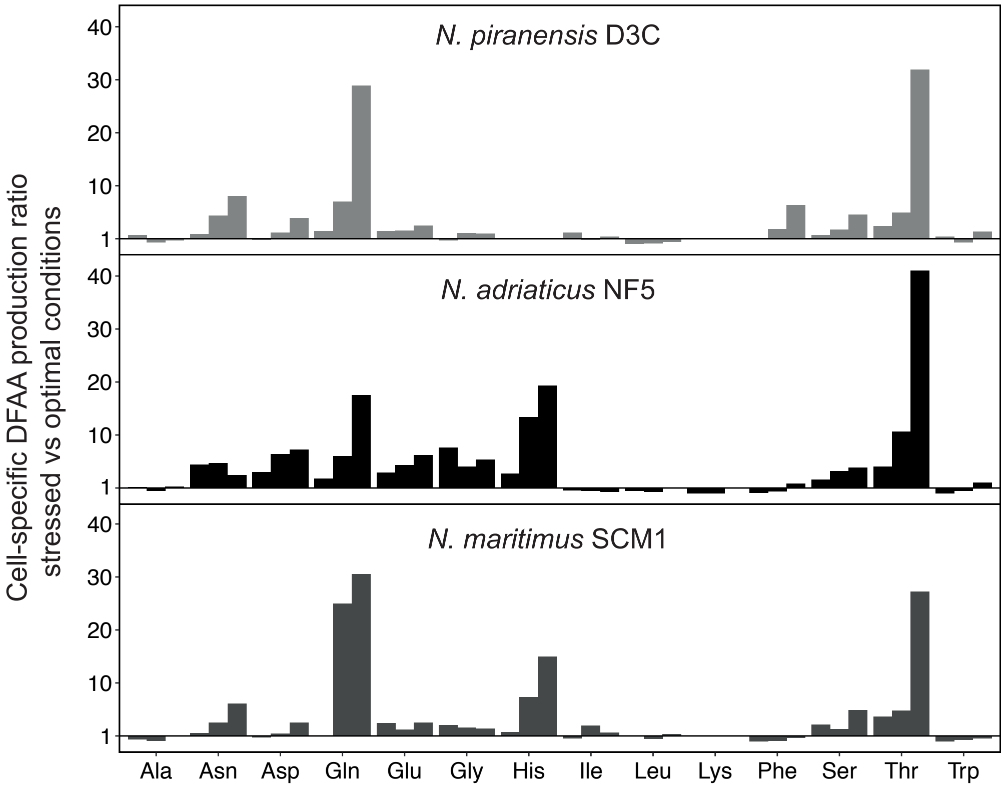


**Figure S3** Mean individual cell-specific DFAA production ratios of stressed vs non-stressed *Nitrosopumilus* strains (amol cell$^{-1}$: amol cell$^{-1}$) at different time points during early to late exponential growth phase (bars from left to right). Values above 1 indicate higher cell-specific production under stress conditions, whereas values below 1 indicate higher cell-specific DFAA production under optimal conditions.

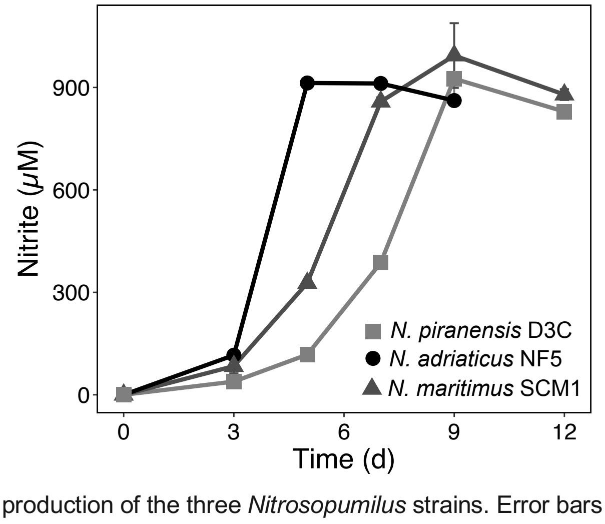


**Figure S4** Nitrite production of the three *Nitrosopumilus* strains. Error bars represent standard deviations of measurements of triplicate cultures.

**Table S1.** Elution gradient applied for the separation of dissolved free primary amino acids by HPLC. A: 89.8 % Methanol, 10 % Milli-Q water, 0.2 % Trifluoroacetic acid; B: Tetrahydrofuran; C: 40 mM $NaH_2PO_4$, pH 7.8

| Elution time [min] | Eluent [%] | | |
|---|---|---|---|
| | A | B | C |
| 0 | 10 | 0 | 90 |
| 2 | 7.5 | 2.5 | 90 |
| 40 | 30 | 2 | 68 |
| 42 | 35 | 1 | 64 |
| 75 | 65 | 0 | 35 |
| 77 | 100 | 0 | 0 |
| 80 | 10 | 0 | 90 |

**Table S2**. Parameters for validation of the HPLC method used to measure dissolved free amino acid (DFAA) concentrations in the presence of high ammonium concentrations. ‡ The limits of detection (LOD) and quantification (LOQ) were determined in Milli-Q, due to presence of background concentrations of amino acids in the culture medium test samples (see Supplementary Material and Methods).

| Substance | Linearity [nM] | $R^2$ | LOD‡ [nM] | LOQ‡ [nM] | Recovery in MQ (n = 5; %) [nM] | | | Recovery in culture medium (n = 5; %) [nM] | | |
|---|---|---|---|---|---|---|---|---|---|---|
| | | | | | 5 | 50 | 500 | 5 | 50 | 500 |
| Aspartic acid | 1-1000 | 0.997-1.000 | ≤ 1 | < 5 | 92.1±8.0 | 97.4±3.6 | 99.0±4.3 | 93.4±2.8 | 83.9±5.2 | 86.2±5.5 |
| Glutamic acid | 1-1000 | 0.997-1.000 | < 1 | < 1 | 100.5±.3.7 | 94.9±2.6 | 94.3±2.3 | 86.4±2.2 | 72.9±3.7 | 73.3±4.3 |
| Asparagine | 1-100 | 0.996-1.000 | < 1 | < 1 | 96.1±.6.1 | 94.5±3.7 | 92.7±3.1 | 116.8±1.2 | 103.1±2.0 | 100.6±3.4 |
| Serine | 1-1000 | 0.998-1.000 | ≤ 1 | ≤ 5 | 107.5±3.5 | 108.3±3.4 | 106.3±2.0 | 66.7±2.3 | 72.1±3.8 | 78.5±2.0 |
| Glutamine | 1-1000 | 0.998-1.000 | < 1 | < 1 | 110.7±4.7 | 108.1±4.2 | 105.8±2.5 | | 91.6±3.27 | 99.1±5.3 |
| Histidine | 1-1000 | 0.999 | < 1 | < 5 | 98.0±8.5 | 98.8±5.6 | 100.6±2.8 | | 119.0±1.1 | 111.6±2.4 |
| Glycine | 1-1000 | 0.998-0.999 | ≤ 1 | < 5 | 103.9±1.4 | 107.0±1.0 | 107.2±3.2 | | 103.2±0.8 | 95.9±0.1 |
| Threonine | 1-1000 | 0.999 | < 1 | ≤ 5 | 106.3±2.8 | 102.8±6.0 | 103.0±3.1 | 79.5±8.6 | 116.5±1.0 | 87.3±1.8 |
| Arginine | 1-1000 | 0.997-0.999 | < 1 | 5 | 93.5±8.7 | 106.9±2.3 | 105.7±4.8 | | 112.4±1.5 | 94.6±2.6 |
| Alanine | 1-1000 | 0.997-0.999 | ≤ 5 | 10 | 109.0±1.3 | 107.3±2.9 | 106.3±2.0 | | 84.7±2.9 | 77.7±4.2 |
| Tyrosine | 1-1000 | 0.996-1.000 | < 1 | < 5 | 111.3±4.3 | 107.3±3.6 | 106.9±2.3 | 84.4±3.5 | 76.8±2.9 | 82.9±4.2 |
| Valine | 1-1000 | 0.993-1.000 | 1 | < 5 | 110.2±2.1 | 106.9±3.2 | 104.7±2.8 | | 69.0±4.3 | 68.8±5.3 |
| Methionine | 5-1000 | 0.989-1.000 | ≤ 1 | ≤ 5 | 115.4±5.2 | 115.1±3.0 | 111.5±3.2 | | 109.7±3.6 | 105.0±2.0 |
| Tryptophan | 1-500 | 0.993-1.000 | ≤ 1 | ≤ 5 | 108.5±4.0 | 107.6±2.1 | 107.0±1.4 | 102.3±4.3 | 85.3±3.1 | 87.3±3.3 |
| Phenylalanine | 1-1000 | 0.991-1.000 | < 1 | < 1 | 110.7±2.6 | 108.0±3.3 | 106.6±2.0 | | 82.4±0.4 | 80.6±4.2 |
| Isoleucine | 1-1000 | 0.995-1.000 | < 5 | < 5 | 109.3±3.3 | 106.3±3.0 | 105.9±2.3 | | 59.5±5.3 | 69.6±6.3 |
| Leucine | 1-1000 | 0.995-1.000 | < 1 | < 5 | 111.3±1.4 | 111.0±2.8 | 107.9±1.3 | 76.8±4.5 | 67.3±3.1 | 73.4±4.3 |
| Lysine | 10-1000 | 0.988-0.999 | ≤ 1 | < 50 | | 97.4±7.5 | 101.9±2.2 | | 72.2±4.9 | 70.6±1.1 |

**Table S3**. Mean dissolved free amino acid (DFAA) concentrations (nM) produced by three *Nitrosopumilus* strains (*N. piranensis* D3C, *N. adriaticus* NF5 and *N. maritimus* SCM1). Standard deviations of triplicate measurements are shown in parentheses. CAT, Catalase addition; bd, below detection; na, not available

| Strain | CAT | Time (d) | Cells x$10^6$ ($ml^{-1}$) | Asp | Glu | Asn | Ser | Gln | His | Gly | Thr | Arg | Ala | Val# | Met# | Trp | Phe | Ile | Leu | Lys |
|---|---|---|---|---|---|---|---|---|---|---|---|---|---|---|---|---|---|---|---|---|
| NF5 | - | 0 | 3.1 | | | | | | | | | | | | | | | | | |
| NF5 | - | 3 | 6.4 | 1 (1) | 4 (1) | 2 (0) | 4 (1) | 4 (0) | 12 (4) | 8 (4) | 23 (12) | bd | bd | na | na | bd | bd | bd | bd | bd |
| NF5 | - | 5 | 6.8 | 2 (0) | 5 (1) | 3 (0) | 11 (6) | 9 (2) | 17 (1) | 35 (2) | 37 (10) | bd | 5 (3) | na | na | bd | bd | 6 (9) | bd | bd |
| NF5 | - | 7 | 6.4 | 2 (0) | 7 (1) | 2 (0) | 17 (7) | 18 (9) | 18 (3) | 64 (6) | 36 (3) | bd | 9 (5) | na | na | bd | 6 (1) | 7 (3) | 3 (0) | 2 (2) |
| NF5 | - | 9 | 6.8 | 3 (1) | 12 (1) | bd | 18 (9) | 16 (2) | 20 (2) | 86 (10) | 37 (2) | bd | 11 (7) | na | na | bd | 9 (2) | 10 (1) | 6 (1) | 8 (1) |
| NF5 | + | 0 | 3.1 | | | | | | | | | | | | | | | | | |
| NF5 | + | 3 | 10.7 | bd | 2 (0) | bd | 2 (1) | 2 (0) | 5 (2) | 2 (1) | 8 (7) | bd | bd | na | na | bd | bd | 5 (5) | 2 (2) | bd |
| NF5 | + | 5 | 23.5 | 1 (1) | 4 (0) | 2 (0) | 9 (2) | 4 (0) | 4 (0) | 25 (2) | 11 (1) | bd | 34 (9) | na | na | bd | 13 (4) | 10 (3) | 15 (2) | 4 (1) |
| NF5 | + | 7 | 31.1 | 1 (1) | 5 (0) | 3 (0) | 17 (6) | 5 (0) | 4 (2) | 49 (5) | 2 (2) | 3 (3) | 34 (4) | 45 (3) | bd | bd | 15 (3) | 4 (1) | 13 (2) | 10 (5) |
| NF5 | + | 9 | 34.6 | 2 (1) | 6 (1) | 2 (0) | 20 (7) | 5 (1) | 6 (3) | 74 (6) | 4 (3) | 4 (3) | 33 (4) | 44 (2) | 2 (0) | 3 (1) | 15 (1) | 4 (1) | 12 (1) | 24 (5) |
| D3C | - | 0 | 2.1 | | | | | | | | | | | | | | | | | |
| D3C | - | 3 | 3.5 | 1 (0) | 3 (1) | 5 (3) | 1 (1) | 4 (1) | bd | 2 (1) | 5 (2) | bd | 4 (3) | na | na | bd | 9 (4) | 3 (2) | bd | na |
| D3C | - | 6 | 4.4 | 2 (0) | 7 (0) | 20 (3) | 2 (1) | 22 (5) | bd | 16 (0) | 20 (2) | bd | 5 (2) | na | na | bd | 17 (3) | 2 (2) | bd | na |
| D3C | - | 9 | 4.3 | 4 (1) | 9 (2) | 29 (6) | 5 (1) | 38 (7) | bd | 35 (3) | 23 (5) | bd | 8 (1) | na | na | 4 (2) | 27 (6) | 15 (6) | bd | na |
| D3C | + | 0 | 2.1 | | | | | | | | | | | | | | | | | |
| D3C | + | 3 | 5 | 2 (2) | 5 (6) | 3 (1) | 2 (3) | 2 (2) | bd | 5 (6) | 2 (3) | bd | 3 (3) | na | na | bd | 16 (11) | 2 (3) | 2 (2) | na |
| D3C | + | 6 | 11.7 | 2 (0) | 7 (1) | 10 (2) | 2 (0) | 7 (2) | bd | 21 (1) | 9 (0) | bd | 48 (4) | na | na | 7 (5) | 16 (9) | 9 (2) | 16 (1) | na |
| D3C | + | 9 | 29.4 | 5 (1) | 17 (3) | 21 (1) | 6 (2) | 9 (2) | bd | 121 (5) | 5 (4) | bd | 82 (12) | 66 (12) | 42 (21) | 10 (9) | 25 (2) | 17 (5) | 28 (5) | na |
| SCM1 | - | 0 | 1.8 | | | | | | | | | | | | | | | | | |
| SCM1 | - | 3 | 4.1 | bd | 2 (0) | 5 (1) | bd | 6 (3) | 2 (2) | 3 (4) | 12 (3) | bd | bd | na | na | bd | bd | bd | bd | na |
| SCM1 | - | 6 | 10.5 | 1 (1) | 7 (1) | 15 (2) | 3 (1) | 24 (6) | 3 (3) | 22 (2) | 23 (3) | bd | 2 (0) | na | na | bd | bd | 3 (3) | 3 (3) | na |
| SCM1 | - | 8 | 10.4 | 2 (0) | 9 (2) | 24 (1) | 6 (2) | 28 (6) | 3 (3) | 46 (4) | 29 (2) | bd | 9 (5) | na | na | bd | 3 (1) | 5 (4) | 6 (1) | na |
| SCM1 | + | 0 | 1.8 | | | | | | | | | | | | | | | | | |
| SCM1 | + | 3 | 5.2 | 1 (0) | 1 (0) | 4 (1) | 1 (1) | bd | 2 (2) | 1 (2) | 3 (3) | bd | 5 (8) | na | na | bd | bd | bd | bd | na |
| SCM1 | + | 6 | 29.4 | 2 (1) | 9 (2) | 12 (2) | 4 (2) | 3 (2) | 3 (3) | 24 (4) | 11 (0) | 2 (2) | 55 (27) | 50 (2) | 25 (8) | 2 (0) | 17 (9) | 3 (2) | 21 (9) | na |
| SCM1 | + | 8 | 48.2 | 3 (0) | 12 (0) | 16 (1) | 5 (0) | 3 (1) | 5 (1) | 88 (2) | na | 7 (2) | 46 (8) | 56 (3) | 29 (9) | 2 (1) | 22 (3) | 5 (2) | 21 (1) | na |

### Val and Met could only be quantified in the absence of $NH_4^+/NH_3$, corresponding to the end of exponential growth under optimal conditions. Thus, T0 measurements could not be subtracted.

**Table S4.** Comparison of intracellular DFAA of three *Nitrosopumilus* strains (*N. piranensis* D3C, *N. adriaticus* NF5 and *N. maritimus* SCM1) and the average amino acid composition encoded by the genomes. Values are given in mol%.

| Amino acid | Genome average | D3C | NF5 | SCM1 |
|---|---|---|---|---|
| Ala | 5.76 | 5.18 | 6.98 | 6.19 |
| Arg | 3.37 | 2.24 | 5.35 | 3.72 |
| Asn | 5.09 | 2.62 | 2.24 | 2.69 |
| Asp | 6.04 | 6.04 | 2.86 | 5.19 |
| Gln | 3.15 | 9.53 | 3.12 | 7.10 |
| Glu | 7.07 | 40.09 | 39.18 | 36.28 |
| Gly | 6.38 | 1.84 | 1.45 | 4.23 |
| His | 1.78 | 1.14 | 2.29 | 1.12 |
| Ile | 9.10 | 2.97 | 3.87 | 3.40 |
| Leu | 8.60 | 2.61 | 1.74 | 2.76 |
| Lys | 8.64 | 3.56 | 7.32 | 5.82 |
| Met | 2.50 | 0.97 | 0.51 | 1.23 |
| Phe | 4.46 | 1.32 | 1.14 | 1.78 |
| Ser | 7.42 | 4.76 | 5.64 | 6.61 |
| Thr | 5.54 | 6.12 | 4.41 | 4.88 |
| Trp | 0.89 | 0.62 | 0.81 | 0.64 |
| Tyr | 3.06 | 3.42 | 5.99 | 3.02 |
| Val | 6.40 | 4.95 | 5.11 | 3.34 |
| Pro | 3.74 | na | na | na |
| Cys | 1.01 | na | na | na |
